# Genome-wide DNA methylation profiling identifies convergent molecular signatures associated with idiopathic and syndromic forms of autism in post-mortem human brain tissue

**DOI:** 10.1101/394387

**Authors:** Chloe C.Y. Wong, Rebecca G. Smith, Eilis Hannon, Gokul Ramaswami, Neelroop N. Parikshak, Elham Assary, Claire Troakes, Jeremie Poschmann, Leonard C. Schalkwyk, Wenjie Sun, Shyam Prabhakar, Daniel H. Geschwind, Jonathan Mill

## Abstract

Autism spectrum disorder (ASD) encompasses a collection of complex neuropsychiatric disorders characterized by deficits in social functioning, communication and repetitive behavior. Building on recent studies supporting a role for developmentally moderated regulatory genomic variation in the molecular etiology of ASD, we quantified genome-wide patterns of DNA methylation in 233 post-mortem tissues samples isolated from three brain regions (prefrontal cortex, temporal cortex and cerebellum) dissected from 43 ASD patients and 38 non-psychiatric control donors. We identified widespread differences in DNA methylation associated with idiopathic ASD (iASD), with consistent signals in both cortical regions that were distinct to those observed in the cerebellum. Individuals carrying a duplication on chromosome 15q (dup15q), representing a genetically-defined subtype of ASD, were characterized by striking differences in DNA methylation across a discrete domain spanning an imprinted gene cluster within the duplicated region. In addition to the dramatic cis-effects on DNA methylation observed in dup15q carriers, we identified convergent methylomic signatures associated with both iASD and dup15q, reflecting the findings from previous studies of gene expression and H3K27ac. Cortical co-methylation network analysis identified a number of co-methylated modules significantly associated with ASD that are enriched for genomic regions annotated to genes involved in the immune system, synaptic signalling and neuronal regulation. Our study represents the first systematic analysis of DNA methylation associated with ASD across multiple brain regions, providing novel evidence for convergent molecular signatures associated with both idiopathic and syndromic autism.

## Introduction

Autism spectrum disorder (ASD) encompasses a collection of complex neuropsychiatric disorders characterized by deficits in social interactions and understanding, repetitive behavior and interests, and impairments in language and communication development. ASD affects ∼1% of the population and confers severe lifelong disability, contributing significantly to the global burden of disease ^1, 2^. Evidence from neuroimaging, neuropathology, genetic and epidemiological studies has led to the conceptualization of ASD as a neurodevelopmental disorder, with etiological origins before birth ^3, 4^. Quantitative genetic analyses have shown that ASD has a strong heritable component ^5^ with an emerging literature implicating rare single base-pair mutations, chromosomal rearrangements, *de novo* and inherited structural genomic variation, and common (polygenic) risk variants in its pathogenesis ^6-9^. Despite the highly heterogeneous role of genetic variation in ASD, studies of transcriptional ^10, 11^ and regulatory genomic variation ^12^ in post-mortem ASD brain provide evidence for a highly-convergent molecular pathology, with individuals affected genetically-defined subtypes of ASD sharing the core transcriptional signatures observed in idiopathic autism cases.

There is increasing evidence to support a role for non-sequence-based genomic variation in the etiology of neurodevelopmental phenotypes including ASD ^13^. The epigenetic regulation of gene expression in the central nervous system is involved in modulating many core neurobiological and cognitive processes including neurogenesis ^14, 15^, neuronal plasticity ^16^ and memory formation ^17, 18^, and is known to be highly dynamic during human brain development ^19^. The dysregulation of epigenetic mechanisms underlies the symptoms of Rett syndrome and Fragile X syndrome, two disorders with considerable phenotypic overlap with ASD ^20-22^, and epigenetic variation has been recently associated with several neurodevelopmental phenotypes including ASD ^12, 23-31^. Current epigenome-wide association studies (EWAS) of autism have focused primarily on DNA methylation, the best characterized and most stable epigenetic modification that acts to influence gene expression via physical disruption of transcription factor binding and through the attraction of methyl-binding proteins that initiate chromatin compaction and gene silencing ^32^. Despite finding evidence for ASD-associated methylomic variation, however, these analyses have been constrained by the analysis of small sample numbers and limited to the assessment of peripheral tissues or a single brain region 12, 23-29.

In this study, we present results from the most systematic analysis of DNA methylation in ASD brain yet undertaken, quantifying methylomic variation in patients with idiopathic ASD (iASD) in addition to patients with a duplication of chromosome 15q11-13 (“dup15q”), which represents the most frequent cytogenetic abnormality associated with ASD occurring in ∼1% of cases ^33, 34^. From each donor we profiled matched post-mortem tissue from three brain regions - prefrontal cortex (PFC), temporal cortex (TC), and cerebellum (CB) – previously implicated in the pathophysiology of ASD. The frontal and temporal lobes, for example, play a role in social cognition, and animal models of ASD highlight cerebellar dysfunction ^35-37^. We find DNA methylation differences in both groups of iASD and dup15q patients, with consistent patterns of variation seen across the two cortical regions, distinct to those identified in CB. In addition to identifying dramatic *cis*-effects of the dup15q duplication in all three brain regions, we identify a significant overlap with the core methylomic differences observed in idiopathic autism iASD cases, reflecting findings from studies of transcriptional variation ^11^ and histone modifications ^12^.

## Results

### Methodological overview

We quantified DNA methylation across the genome using the Illumina Infinium HumanMethylation450 BeadChip (“450K array”) in 223 post-mortem tissue samples comprising prefrontal cortex (PFC), temporal cortex (TC) and cerebellum dissected from 43 donors with ASD (including 7 patients with dup15q syndrome and 36 patients with idiopathic ASD (iASD)) and 38 non-psychiatric control subjects. After implementing a stringent quality control (QC) pipeline (see **Methods**), we obtained high-quality DNA methylation data from 76 PFC samples (n = 36 iASD patients, n = 7 dup15q patients, n = 33 controls), 77 TC samples (n = 33 iASD patients, n = 6 dup15q patients, n = 38 controls) and 70 CB samples (n = 34 iASD patients, n = 7 dup15q patients, n = 29 controls) (**Supplementary Table 1**). Our primary analyses focused on identifying differentially methylated positions (DMPs) and differentially methylated regions (DMRs) associated with iASD and dup15q, controlling for cellular heterogeneity and other potential confounds, exploring the extent to which signals were shared across idiopathic and syndromic autism cases. Finally, we employed weighted gene co-methylation network analysis (WGCNA) to undertake a systems-level view of the DNA methylation differences associated with both iASD and dup15q across the three brain regions. An overview of our experimental approach is given in **Supplementary Figure 1**.

**Table 1.**
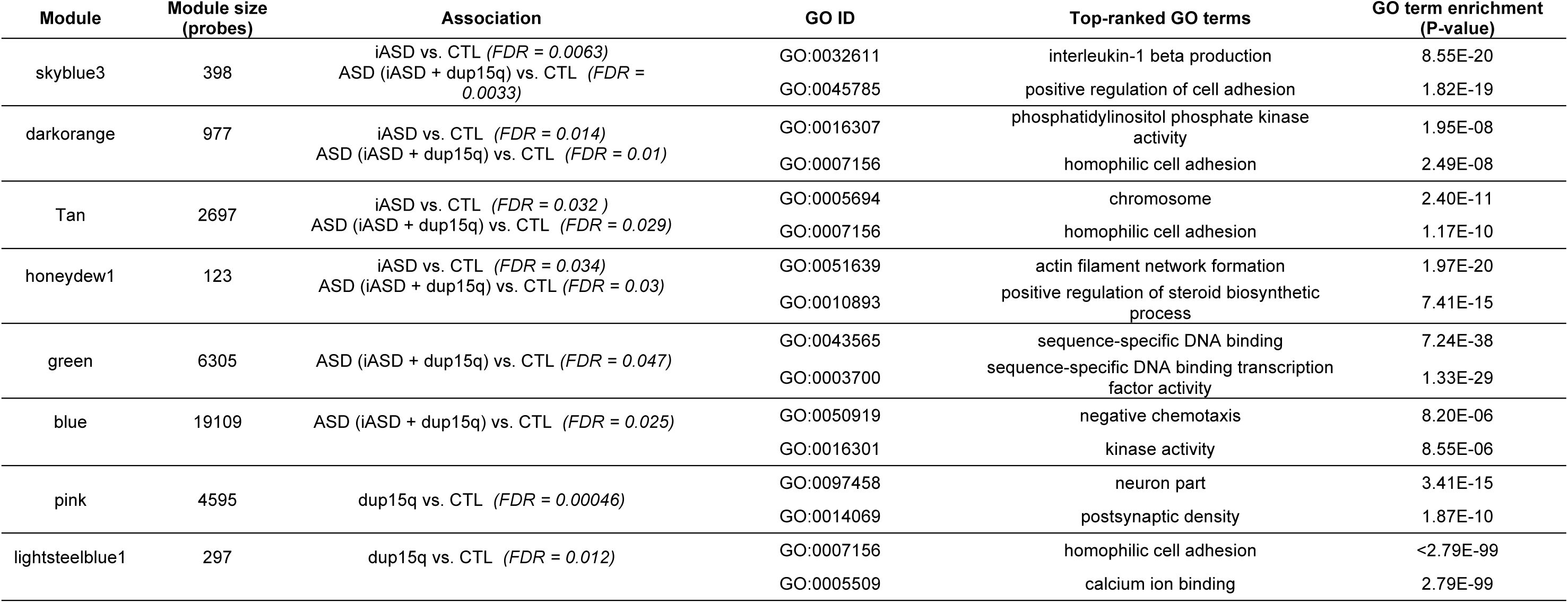
Cortical co-methylation modules associated with ASD. A full list of GO pathways enriched amongst genes annotated to the Illumina 450K probes in each module is given in **Supplementary Table 17**.

**Figure 1.**
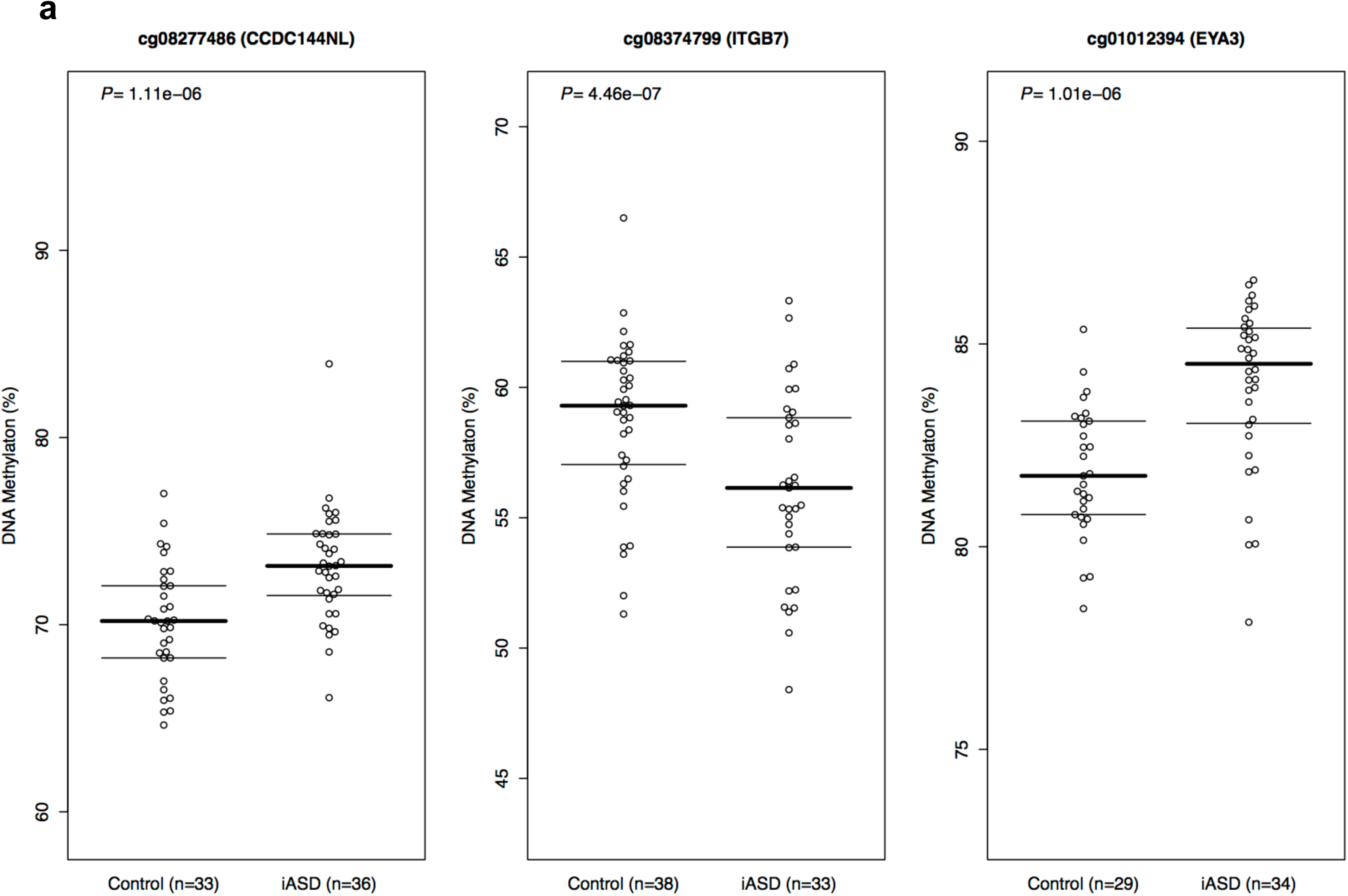
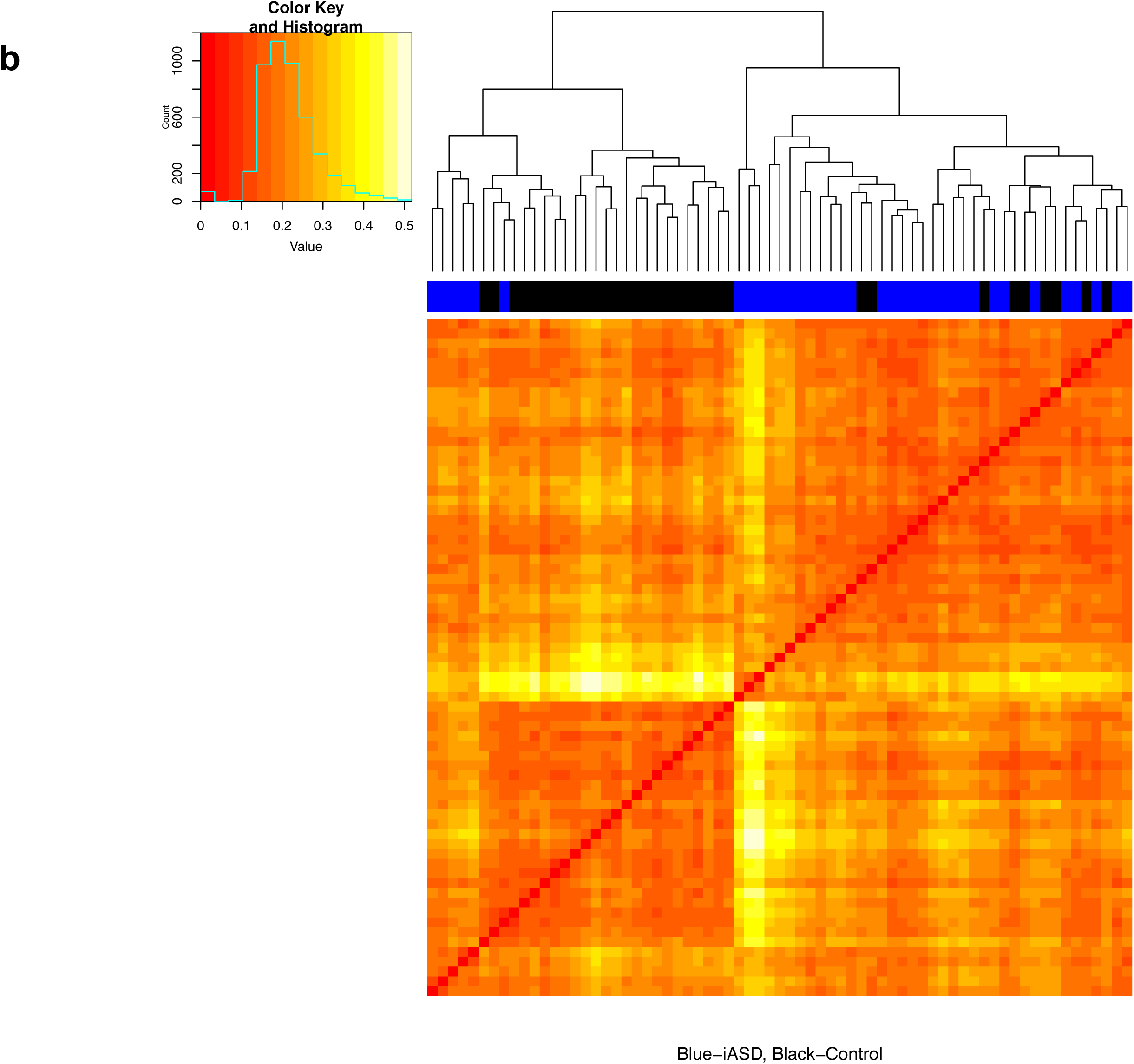
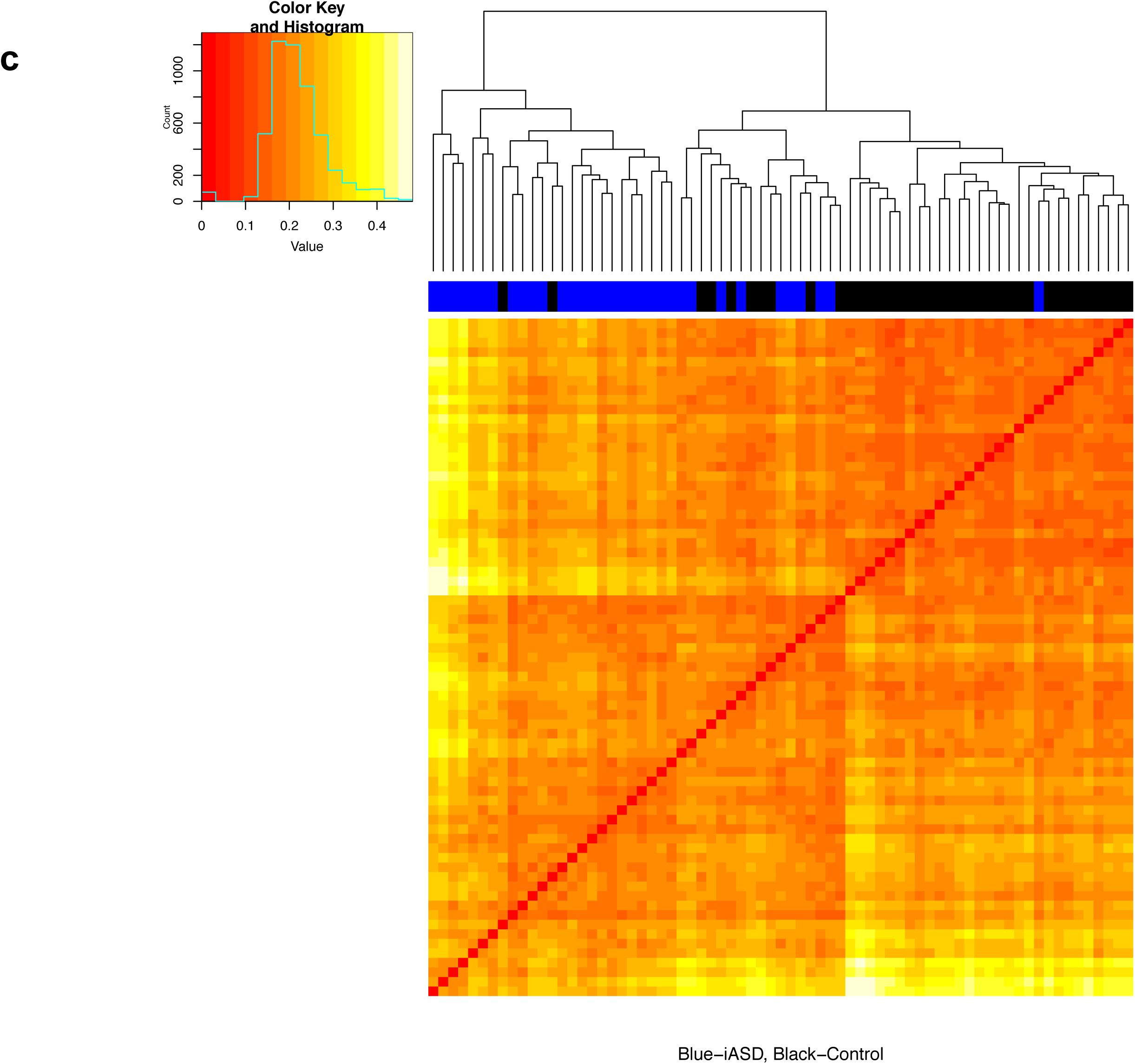
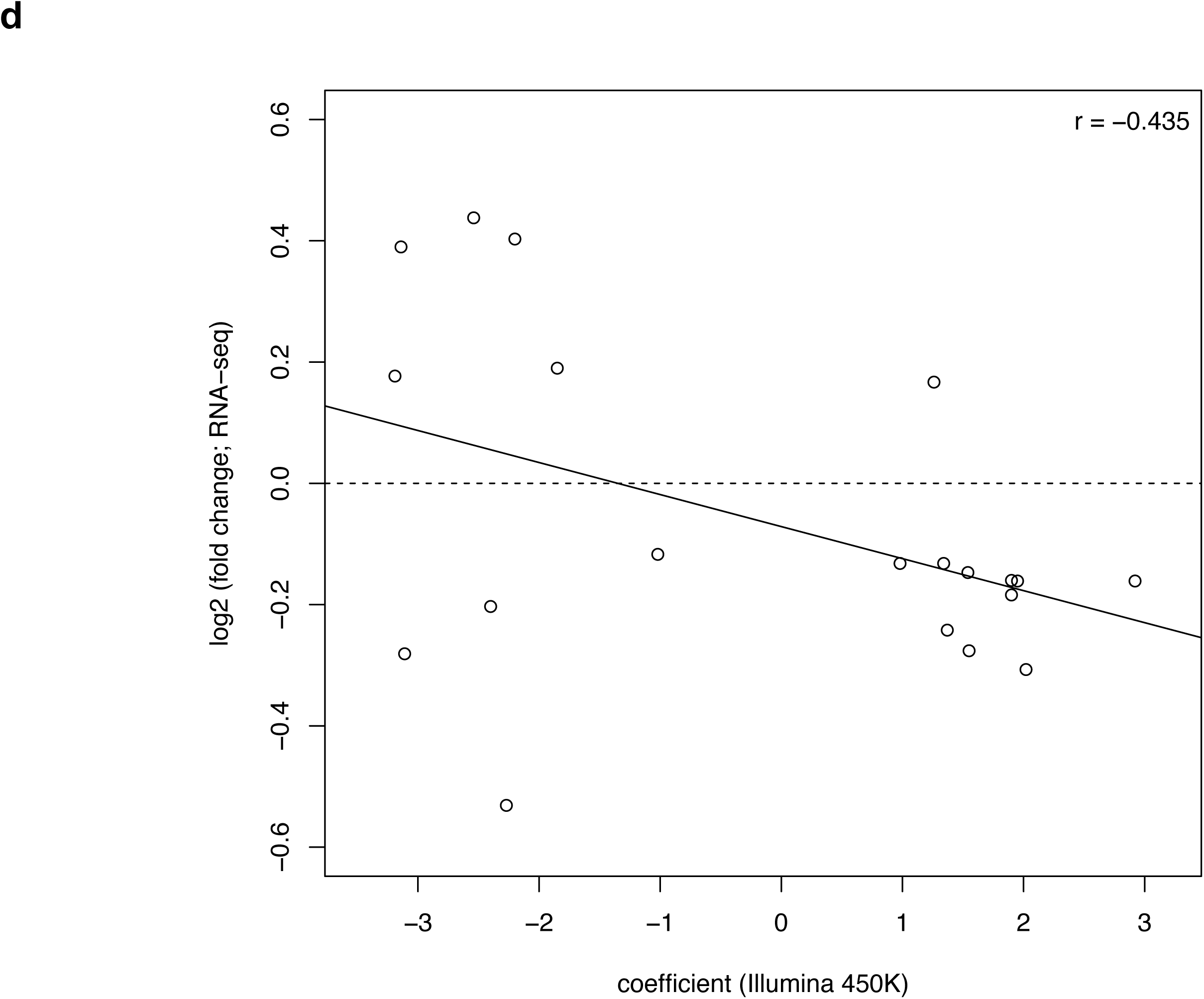
DNA methylation differences at sites associated with idiopathic autism cluster cases and controls and are correlated with gene expression differences. Site-specific changes in DNA methylation associated with idiopathic autism. **(a)** Shown are the top-ranked iASD-associated differentially methylated positions in prefrontal cortex (cg08277486), temporal cortex (cg08374799), and cerebellum (cg01012394). A complete list of all iASD-associated DMPs (P < 5e-05) is provided in **Supplementary Tables 2-4. (b-c)** Hierarchical clustering of samples based on DNA methylation at iASD-associated DMPs in the cortex. Shown is the clustering of samples based on DNA methylation levels (red = low, yellow = high) at iASD-associated DMPs (P < 5e-05) in **(b)** prefrontal cortex and **(c)** temporal cortex. In both cortical regions, the two primary clusters are clearly aligned to disease status (prefrontal cortex: cluster 1 = 77% iASD, cluster 2 = 20% iASD; temporal cortex: cluster 1 = 92% iASD, cluster 2 = 22% iASD)). **(d)** Effect sizes at iASD-associated DMPs are negatively correlated with gene expression level. Shown is the relationship between DNA methylation difference (X-axis) and gene expression difference (Y-axis) for genes annotated to cortical DMPs identified in a RNA-seq study performed on an overlapping set of samples.

### DNA methylation differences between iASD cases and controls are consistent across cortical regions

No global differences in DNA methylation - estimated by averaging across all Illumina 450K array probes (n=417,460) included in our analysis - were identified between iASD patients and control subjects in any of the three brain regions (PFC: iASD = 48.4%, controls = 48.5%; TC: iASD = 48.4%, controls = 48.4%; CB: iASD = 46.4%, controls = 46.4%). We observed a robust positive correlation between the estimated ‘DNA methylation age’ - calculated using an epigenetic clock based on DNA methylation values ^38, 39^ - and recorded chronological age for each of the brain regions (PFC: r = 0.98, TC: r = 0.97. CB: r = 0.94) (**Supplementary Figure 2**), with no evidence for differential ‘epigenetic aging’ in iASD patients (PFC: P = 0.10, TC: P = 0.24, CB: P = 0.80). These findings indicate that ASD is not associated with any systemic differences in DNA methylation across the probes included on the Illumina 450K array in the brain regions tested, reflecting findings in studies of other complex neuropsychiatric phenotypes including Alzheimer disease ^40^ and schizophrenia ^41^.

**Figure 2.**
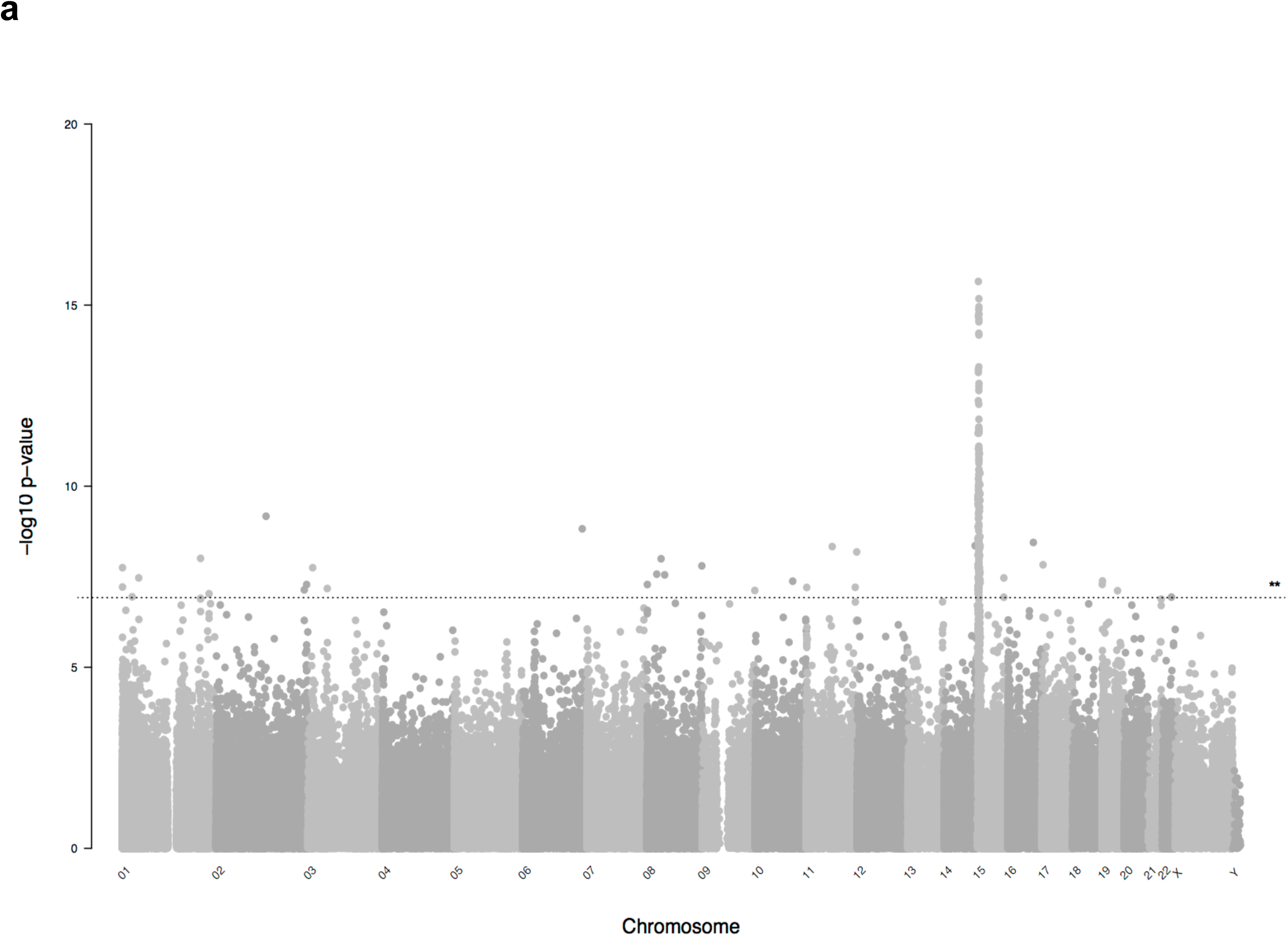
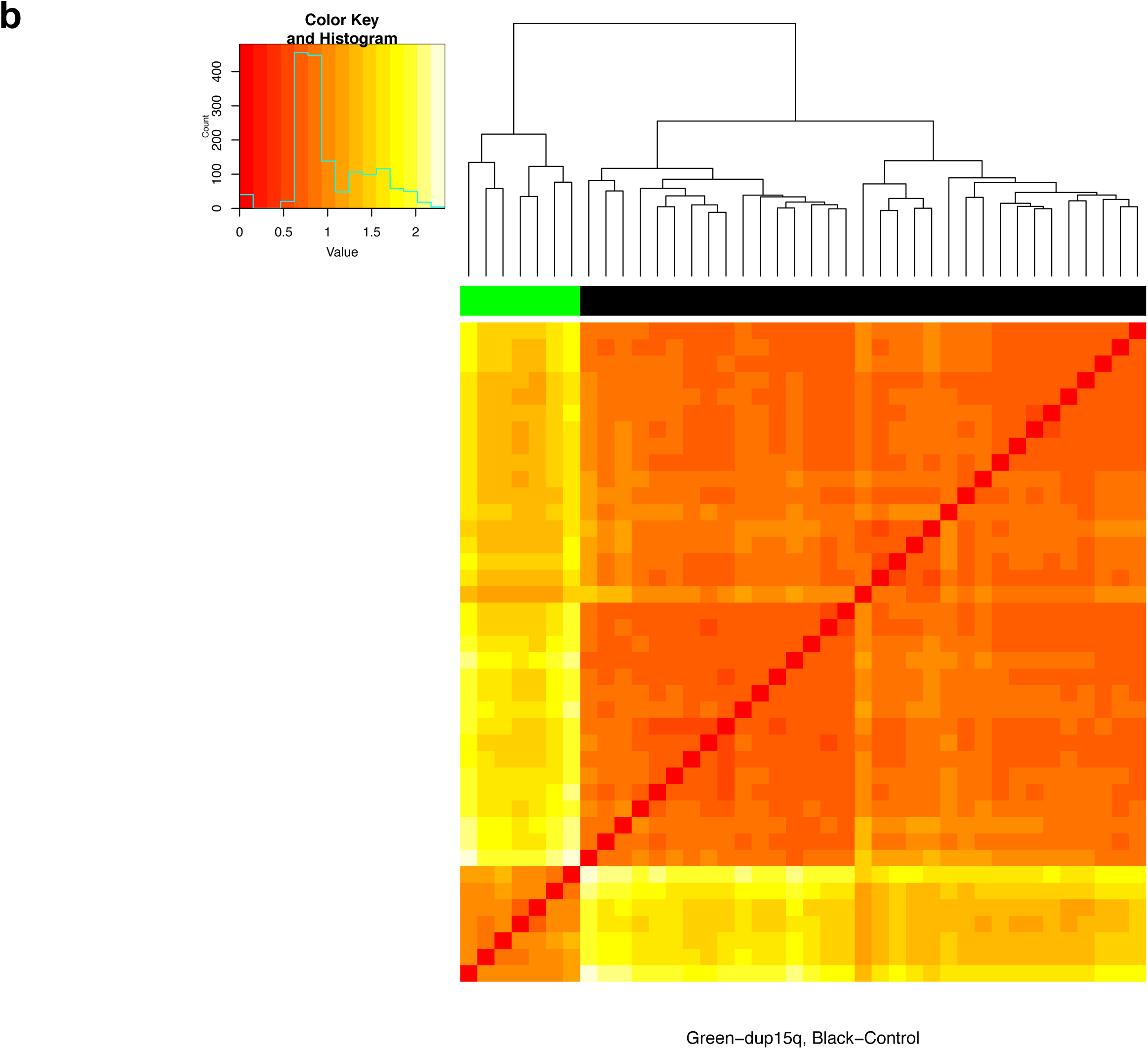
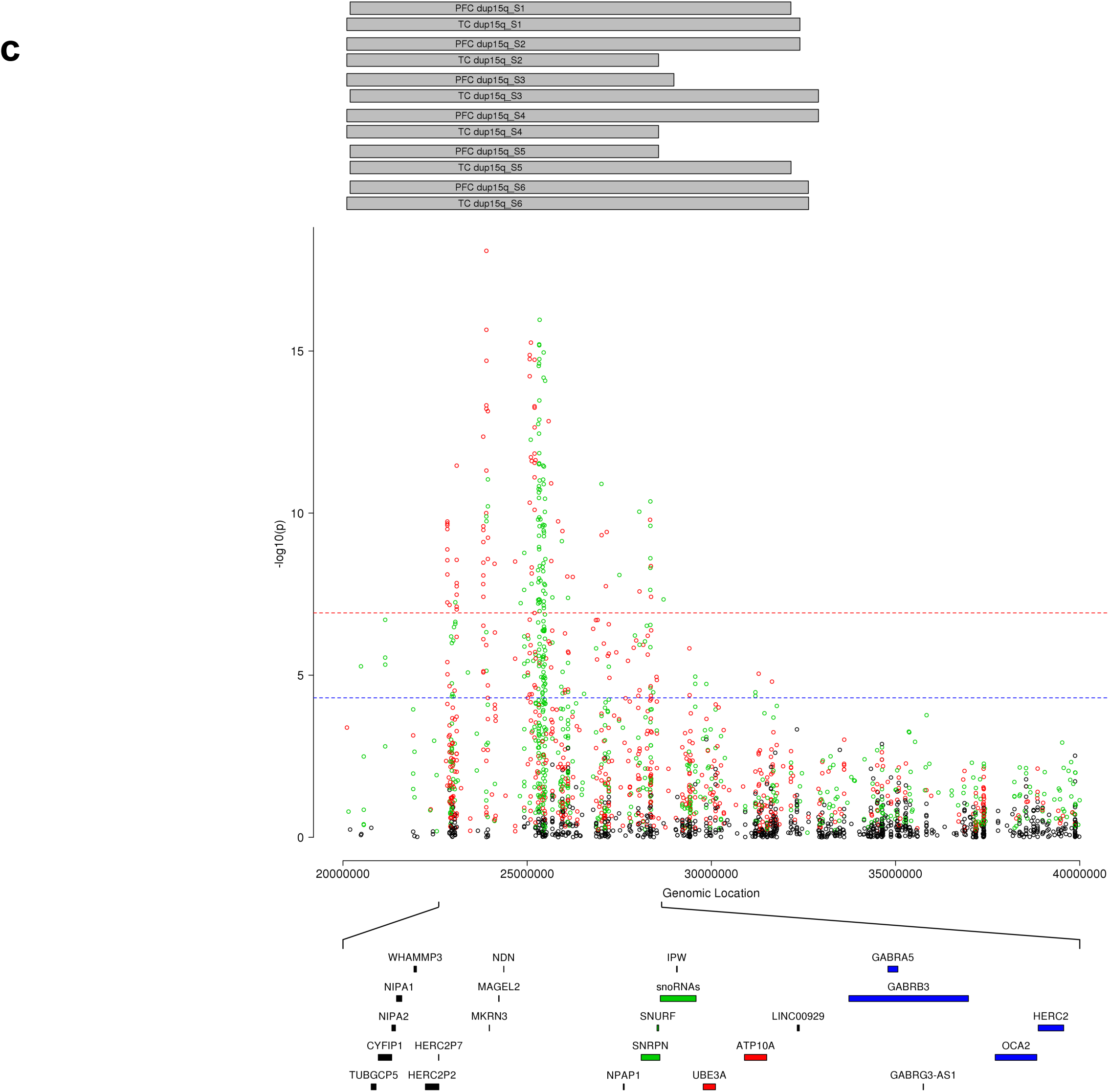
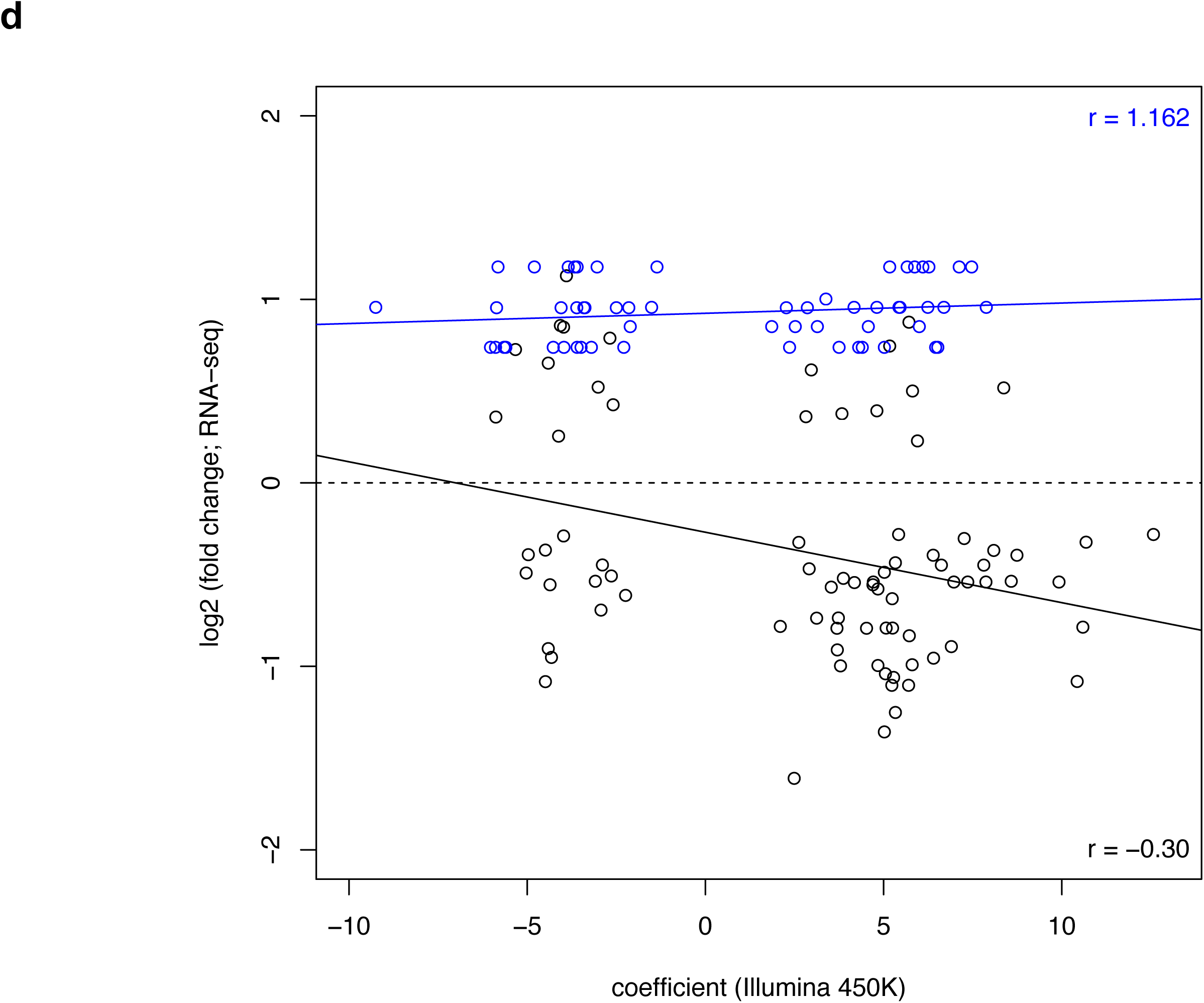
The duplication of chromosome 15q is associated with distinct DNA methylation profiles in all three brain regions. **(a)** Manhattan plot of P-values from a multi-level model used to identify consistent dup15q-associated differences across both cortical regions (PFC and TC). The majority of dup15q-associated DMPs are located within the duplicated region. *The line highlights a bonferroni significance threshold of P < 1.198X10^−7^. **(b)** Hierarchical clustering of samples based on DNA methylation at dup15q-associated DMPs in the cortex. Shown is the clustering of samples based on PFC DNA methylation levels (red = low, yellow = high) at dup15q-associated DMPs (P < 5e-05). **(c)** Cortical dup15q-associated DMPs are localized within a narrow region within the duplicated region. Shown is the distribution of P-values across the dup15q region from our cross-cortex (FC and TC) model. Differentially methylated positions are stratified by direction of effect (red = hypermethylated in dup15q ASD, green = hypomethylated in dup15q ASD). Shown at the top are the estimated break-points for individual dup15q samples derived from PFC and TC DNA methylation data for each individual donor. The dup15q differentially methylated domain includes clusters of probes that are both hyper- and hypo-methylated overlapping a known imprinted gene cluster containing paternally-expressed (green), maternally-expressed (red) and biallelically-expressed (blue) genes. **(d)** Effect sizes at dup15q-associated DMPs are inversely correlated with gene expression level. Shown is the relationship between DNA methylation difference (X-axis) and gene expression difference (Y-axis) for genes annotated to cortical DMPs identified in a RNA-seq study performed on an overlapping set of samples. The overall correlation between DNA methylation is gene expression was −0.313, although there were striking differences between loci within the dup15q region (r = 0.162; colored in blue) and elsewhere in the genome (r = −0.300; colored in black).

We next used a linear model including covariates for sex, age, brain-bank and neuronal cell proportions derived from the DNA methylation data (except in the CB, as described in the **Methods**) to identify iASD-associated differentially methylated positions (DMPs) across the genome in each of the three brain regions. The top ranked iASD-associated DMPs in each brain region (PFC: cg08277486, which is located within *CCDC144NL* and hypermethylated in patients compared to controls (P = 1.11e-06); TC: cg08374799, which is located immediately upstream of *ITGB7* and hypomethylated in patients compared to controls (P = 4.46e-07); CB: cg01012394, which is located immediately upstream of *EYA3* and hypermethylated in patients compared to controls (P = 1.01e-06)) are shown in **Figure 1a**, with a list of all DMPs (P < 5e-05) detailed in **Supplementary Tables 2-4**. Of note, iASD-associated DNA methylation differences are considerably more pronounced in both cortical regions (PFC: n = 31 DMPs; TC: n = 52 DMPs) than the cerebellum (n = 2 DMPs). Hierarchical clustering of samples based on DNA methylation levels at these cortical DMPs distinguishes relatively well between iASD cases and controls in both PFC (**Figure 1b**) and TC (**Figure 1c**). As reported in previous analyses of epigenetic variation in the human brain ^41, 42^, our data show that - at a global level - the patterns of DNA methylation in the CB are very distinct to the two cortical brain regions included in this study (**Supplementary Figure 3**). Effect sizes at iASD-associated DMPs are highly correlated between the two cortical regions, but not between cortex and CB [top 100 PFC DMPs (PFC vs TC: r = 0.77, P = 3.06e-21; PFC vs CB: r = 0.14, P = 0.18), top 100 TC DMPs (TC vs PFC: r = 0.81, P = 2.48e-24; r = 0.17, P = 0.09) and top 100 CB DMPs (CB vs PFC: r = 0.005, P = 0.96; CB vs TC: r = −0.03, P = 0.77)] (**Supplementary Figure 4**). Given the striking consistency of effects across cortical regions we used a multi-level linear mixed model (see **Methods**) to maximize our power to identify consistent iASD-associated differences across PFC and TC (**Supplementary Figure 5** and **Supplementary Table 5**). We identified 157 DMPs (P < 5e-05), with the top-ranked cross-cortex iASD-associated difference (cg14392966, P = 1.77E-08) being located in the promoter region of *PUS3* (+6bp) and upstream of *DDX25* (+1136bp) on chromosome 11q24.2. Using data from an RNA-seq analysis of an overlapping set of samples ^11^, we explored the extent to which genes annotated to these DMPs were also differentially expressed in iASD cortex. Of the 111 DMPs annotated to a gene, 20 (18.0%) were annotated to a transcript found to be differentially expressed in iASD cortex (FDR < 0.1) (**Supplementary Table 6**), with an overall negative correlation (r = −0.435) between DNA methylation and gene expression effects sizes across these probe-gene pairs (**Figure 1d**).

**Figure 3.**
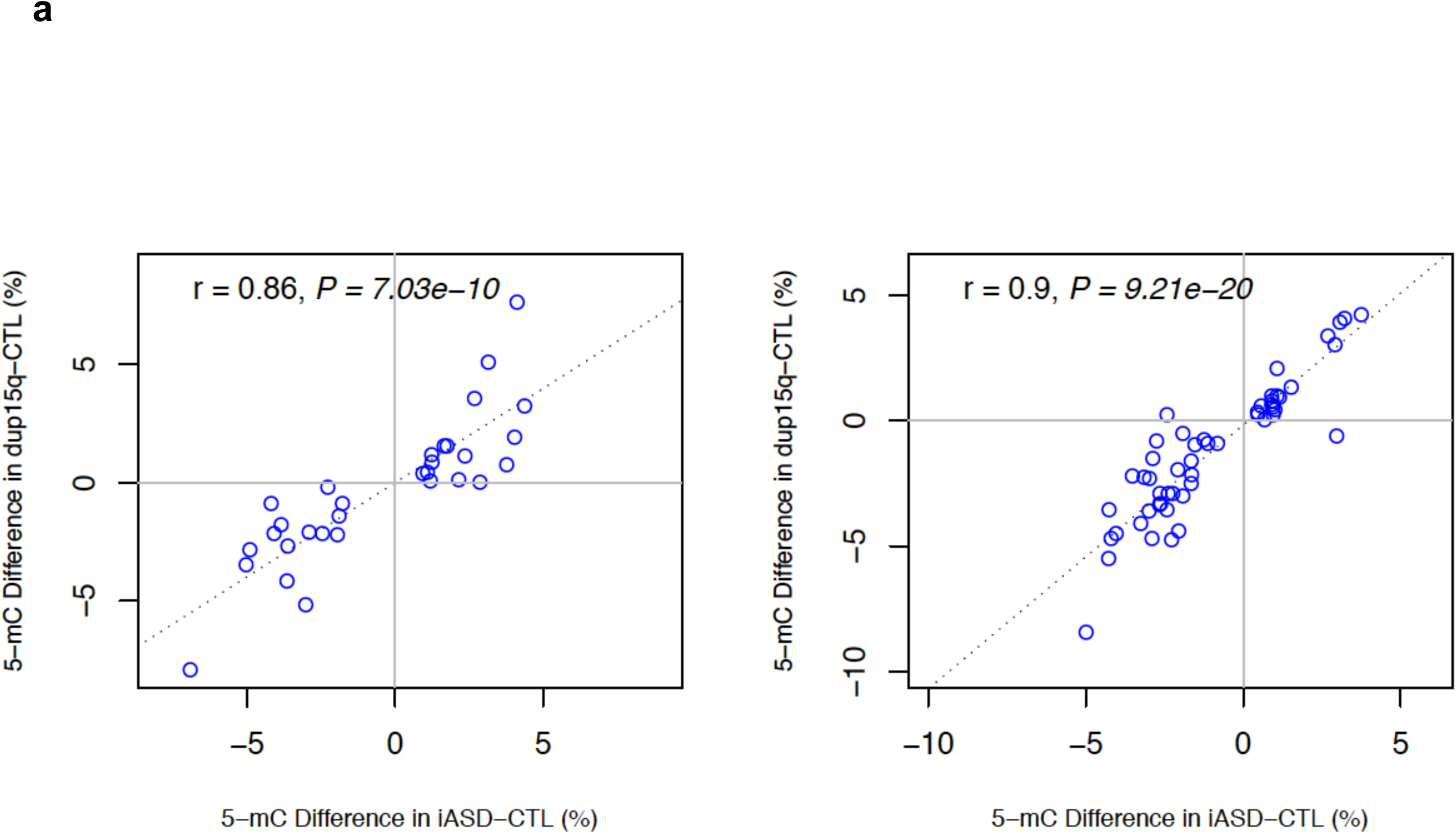
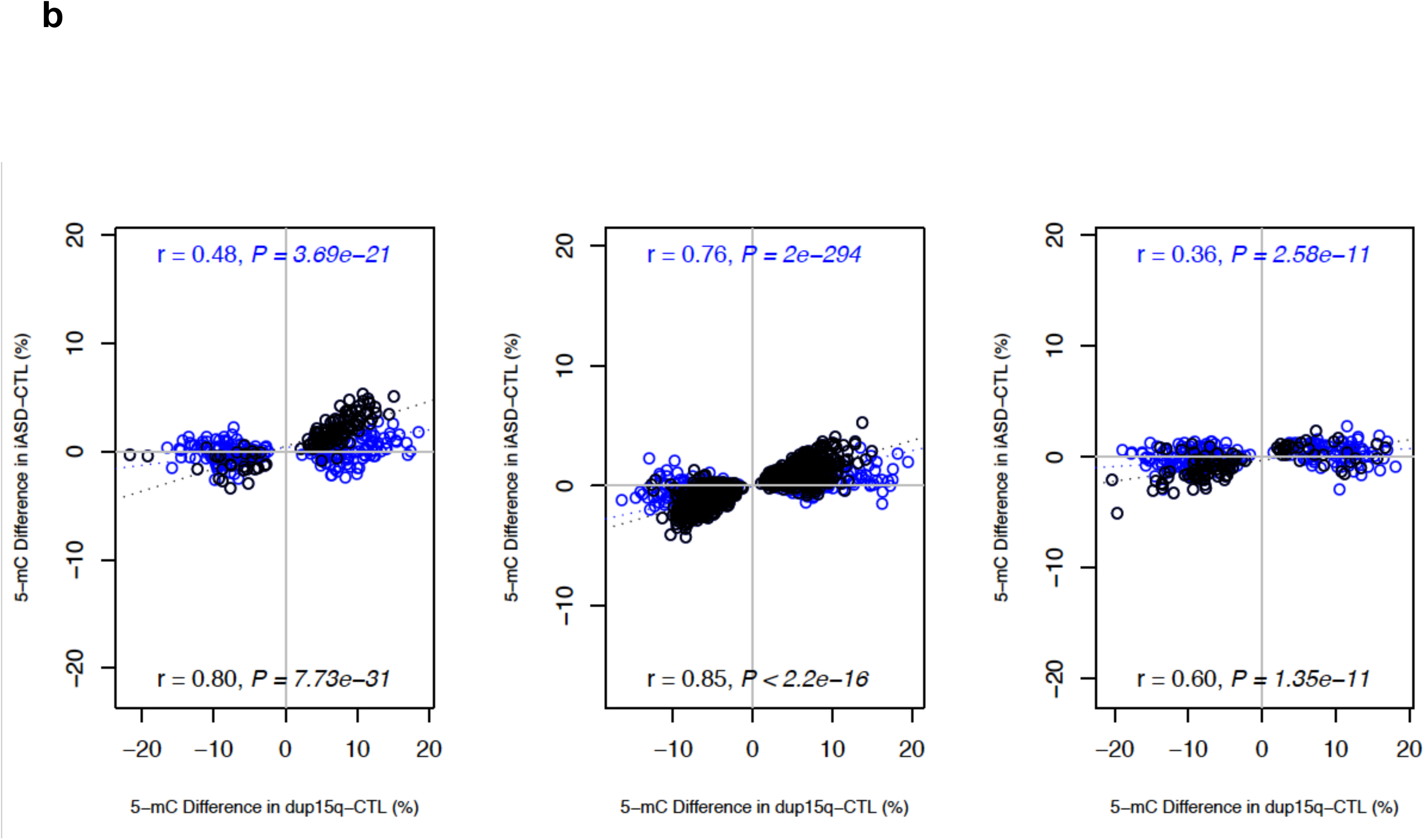
Consistent DNA methylation differences are seen in iASD and dup15q patients. **(a)** Effect sizes at cortical iASD-associated DMPs are significantly correlated between iASD and dup15q patients. Shown is the correlation in effect sizes for PFC (r = 0.86, P = 7.03e-10, left panel) and TC (r = 0.9, P = 9.21e-20, right panel). **(b)** Effect sizes at dup15q-associated DMPs are highly correlated between dup15q and iASD patients. Shown is the correlation in effect sizes for PFC (across all probes r= 0.48, P = 3.69e-21; probes outside of the dup15q region r = 0.80, P = 7.73e-31; left panel), TC (across all probes r = 0.76, P = 2e-294; probes outside of the dup15q region r = 0.85, P < 2.2e-16; middle panel), and CB (across all probes r = 0.36, P = 2.58e-11; probes outside of the dup15q region r = 0.60, P = 1.35e-11; right panel).

### Dup15q, a genetically-defined subtype of ASD, is associated with striking differences in DNA methylation across an imprinted gene cluster within the duplicated region

The duplication of human chromosome 15q11-13 (“dup15q”) is the most frequent cytogenetic abnormality associated with ASD, occurring in ∼1% of cases ^33, 34^. In addition to the iASD cases profiled in this study, we quantified DNA methylation in PFC, TC, and CB tissue from seven individuals with dup15q; using an established method to identify copy number variation (CNV) from Illumina 450K data ^43^, we confirmed dup15q status in each of the brain regions profiled in the seven carriers (**Supplementary Figures 6-8).** As with the iASD patients, no global differences in DNA methylation were identified between dup15q carriers and controls in any of the three brain regions (PFC: dup15q carriers = 48.5%, controls = 48.5%; TC: dup15q carriers = 48.4%, controls = 48.4%; CB: dup15q carriers = 46.4%, controls = 46.4%). Again, we observed a strong positive correlation between the estimated ‘DNA methylation age’ with recorded chronological age for each of the brain regions (**Supplementary Figure 2**). Although there was no evidence for differential ‘epigenetic aging’ in dup15q patients compared to controls in either cortical region (PFC: P = 0.06, TC: P = 0.45), there was a nominally-significant association in CB (P = 0.012) with dup15q carriers characterized by decelerated epigenetic aging compared to controls. Overall, these findings indicate that like iASD patients, dup15q carriers are not characterized by systemic differences in DNA methylation across the probes included on the Illumina 450K array in the brain regions tested.

Using a linear model including covariates for sex, age, brain-bank and neuronal cell proportions derived from the DNA methylation data (except in the CB as described in the **Methods**) we identified numerous DMPs in dup15q carriers in each of the three brain regions (**Supplementary Tables 7-9** and **Supplementary Figures 9-11**). As in our analysis of iASD, we used a multi-level model (see **Methods**) to identify consistent dup15q-associated differences across PFC and TC (**Figure 2** and **Supplementary Table 10**). Using *comb-p* ^44^ we also identified spatially correlated regions of differential DNA methylation associated with dup15q status (Sidak corrected P < 0.05) (**Supplementary Tables 11-14**). Our analyses revealed striking cis-effects on DNA methylation, with the majority of significant DMPs (**Figure 2c**, **Supplementary Figures 6-8**) and DMRs located within a ∼ 7Mb cluster in the 15q11.1-13.2 duplication region. Despite these strong *cis*-effects, however, sites within the 15q duplicated region are not ubiquitously differentially methylated in CNV carriers. Dup15q-associated DMPs were found to be focused in a specific region within the duplication, with this discrete differentially methylated domain includes clusters of probes that are both hyper- and hypo-methylated in carriers (**Figure 2c** and **Supplementary Figures 6-8**). Interestingly, these DMPs overlap a genomically imprinted gene cluster within the duplicated region containing transcripts monoallelically expressed from either the paternal (*SNRPN, snoRNAs*) or maternal (*UBE3A, ATP10A*) alleles. Although DMPs located in the dup15q region are highly consistent across each of the three brain regions, the overall pattern of dup15q-associated variation is more similar between the two cortical regions than between cortex and cerebellum (**Supplementary Figure 12**), reflecting the patterns observed for iASD. Interestingly, despite the large effects observed within the dup15q region, a number of DMPs (**Supplementary Tables 7-10**) and DMRs (**Supplementary Tables 11-14**) outside the vicinity of the duplication were also identified in each of the three brain regions, suggesting that structural variation on chromosome 15 may influence regulatory genomic variation at other chromosomal locations in *trans*. Using data from an RNA-seq analysis of an overlapping set of samples ^11^, we explored the extent to which genes annotated to these DMPs were also differentially expressed in dup15q ASD cortex. Of the 699 DMPs annotated to a gene, 139 (19.9%) were annotated to a transcript found to be differentially expressed in dup15q cortex (FDR < 0.1) (**Supplementary Table 15**), with an overall negative correlation (r = −0.313) between DNA methylation and gene expression effects sizes across these probe-gene pairs (**Figure 2d**). Of note, there were striking differences in the correlation between DNA methylation and gene expression for loci within the dup15q region (r = 0.162) compared to those elsewhere in the genome (r = −0.300).

### Methylomic differences are shared between iASD and dup15q carriers

Building on a recent analysis of gene expression ^11^ that revealed a core pattern of cortical transcriptional dysregulation observed in both iASD and dup15q carriers, we next examined the extent to which disease associated DNA methylation differences are shared between these two distinct subgroups of autism patients. Effect sizes at iADS DMPs (P < 5×10^−5^) were significantly correlated between iASD and dup15q patients (PFC: r = 0.86, P = 7.03e-10; TC: r = 0.9, P = 9.21e-20; CB: not tested because of the small number of significant DMPs) (**Figure 3a)**. Likewise, effect sizes at dup15q DMPs (P < 5e-05) were found to be highly correlated between dup15q and iASD patients (PFC: r= 0.48, P = 3.69e-21; TC: r = 0.76, P = 2e-294; CB: r = 0.36, P = 2.58e-11) (**Figure 3b)**; of note, although these correlations were particularly strong for probes outside of dup15q region (PFC: r = 0.80, P = 7.73e-31; TC: r = 0.85, P < 2.2e-16; CB: r = 0.60, P = 1.35e-11), significant correlations were also seen for dup15q-associated DMPs located in the duplicated region (PFC: r = 0.22, P = 8.4e-4; TC: r = 0.53, P = 8.59e-18; CB: r = 0.22, P = 0.001). Hierarchical clustering of samples based on DNA methylation values at iASD-associated DMPs (P < 5 x10^−5^) shows that dup15q carriers cluster together with iASD cases (**Supplementary Figures 13** and **14)**, highlighting convergent methylomic signatures associated with both idiopathic and syndromic forms of autism.

### Cortical co-methylation modules associated with ASD are enriched for immune, synaptic and neuronal processes

We next used weighted gene co-methylation network analysis (WGCNA) ^45^ to characterize systems-level differences in DNA methylation associated with ASD. We built co-methylation networks using all ‘variable’ DNA methylation sites (defined as those where the range of DNA methylation values for the middle 80% of individuals was greater than 5%; N = 251,311) using cross-cortex (PFC and TC) data from all donors (see **Methods**). WGCNA identified 61 co-methylation modules (**Supplementary Table 16**) and we used the ‘module eigengene’ (i.e. the first principal component) for each module to explore differences between controls (n = 29), individuals with iASD (n = 30), individuals with dup15q (n = 6), and a combined ASD group (n = 36). We identified several co-methylation modules robustly associated (FDR < 0.05) with at least one diagnostic category (**Table 1, Supplementary Figure 15**). We tested whether the genes annotated to probes in each ASD-associated co-methylation module were enriched for specific gene ontology (GO) pathways using a method that groups related pathways to control for the hierarchical structure of the ontological annotations (see **Table 1** and **Supplementary Table 17**), identifying a number of pathways relevant to the known etiology of ASD. For example, the most enriched pathways amongst genes annotated to probes in the “skyblue3” module, which was associated with both iASD (FDR = 0.0063) and the combined ASD group (FDR = 0.0033), are related to immune function (e.g. interleukin-1 beta production, P = 8.55E-20), consistent with findings from genetic ^46^, transcriptomic ^47^, and epidemiological data ^48, 49^. Amongst genes annotated to probes in the “darkorange” module, which was also associated with both iASD (FDR = 0.014) and the combined ASD group (FDR = 0.01), the top-ranked pathways were related to synaptic signaling and regulation, in particular phosphatidylinositol 3-kinase (PI3K) activity (P = 1.95E-08), which regulates synaptic formation and plasticity ^50^. Also of note, pathways enriched amongst genes annotated to probes in the ‘pink’ module, which was associated with dup15q carriers (FDR = 0.00046) were related to pathways important in neurons (P = 3.41E-15) and postsynaptic density (P = 1.87E-10). A full list of all significant pathways for each of the co-methylation modules is given in **Supplementary Table 17**.

## Discussion

In this study, we quantified DNA methylation in 223 post-mortem tissue samples isolated from the prefrontal cortex, temporal cortex and cerebellum dissected from 43 donors with ASD and 38 non-psychiatric control subjects. To our knowledge, this represents the most systematic analysis of DNA methylation in ASD using disease-relevant tissue, and the first to compare variation identified in idiopathic and syndromic forms of ASD. We report ASD-associated DNA methylation differences at numerous CpG sites with more pronounced effects in both cortical regions compared to the cerebellum. This finding is consistent with previous gene expression studies illustrating that ASD-related molecular changes are substantially smaller in the cerebellum compared to the cortex ^11^.

Although structural variation on chromosome 15 was found to be associated with striking cis-effects on DNA methylation, with a discrete differentially methylated domain spanning an imprinted gene cluster within the duplicated region, variation in DNA methylation associated with autism in the cerebral cortex was highly correlated between iASD and dup15q patients. These results suggest that there are convergent molecular signatures in the cortex associated with different forms of ASD, reinforcing the findings from a recent study of gene expression ^11^ and H3K27ac^12^ undertaken in an overlapping set of samples. Co-methylation network analyses highlighted systems-level changes in cortical DNA methylation associated with both iASD and dup15q, with associated modules being enriched for sites annotated to genes involved in the immune system, synaptic signalling and neuronal regulation. Our results corroborate findings from other DNA methylation ^26, 30, 31^ and gene expression analyses ^10, 11^ which have concluded that ASD-related co-methylation and co-expression modules are significantly enriched for synaptic, neuronal and immune dysfunction genes. Finally, we used existing RNA-seq data on an overlapping set of samples to show a remarkable overlap of cortical ASD-associated differentially methylated positions with differential gene expression. Future studies should focus on further understanding the transcriptional consequences of the observed associations, and testing whether these associations are causal or a consequence of disease and/or medication.

This study has several strengths. Our epigenome-wide analysis of ASD is, to our knowledge, the largest post-mortem cohort so far and included tissue from three brain regions that have been previously implicated in the pathophysiology of ASD. This contrasts with previous studies that have been undertaken on much smaller numbers of samples and focused on only one or two brain regions. The inclusion of both idiopathic ASD patients and dup15q carriers in our analyses enabled us to explore evidence for convergent molecular signatures associated with both idiopathic and syndromic forms of autism.

Despite this being the first study to quantify DNA methylation across three different brain regions from both idiopathic and syndromic ASD patients and controls, this study has a number of important limitations that should be considered when interpreting the results. First, DNA methylation was quantified using the Illumina 450K array; although this is a robust and highly reliable platform with content spanning regulatory regions associated with the majority of known annotated genes, it interrogates DNA methylation at a relatively small proportion of sites across the whole genome. Second, because epigenetic processes play an important role in defining cell-type-specific patterns of gene expression ^51, 52^, the use of bulk tissue from each brain region is a potential confounder in DNA methylation studies^53^. Despite our efforts to control for the effect of cell type diversity in DNA methylation quantification in our analyses using *in silico* approaches, this approach is not suitable to estimate the neuronal proportion in the cerebellum and cannot inform us about disease relevant DNA methylation changes specific to individual brain cell types. Of note, our general findings are in line with those reported by Nardone and colleagues^26^ which interrogated DNA methylation in cell-sorted cortical neurons (from 16 ASD cases and 15 controls). Third, there is increasing awareness of the importance of 5-hydroxymethyl cytosine (5-hmC) in the human brain ^54^, although this modification cannot be distinguished from DNA methylation using standard bisulfite-based approaches. It is plausible that many of the ASD-associated differences identified in this study are confounded by modifications other than DNA methylation. To date, no study has evaluated the role of genome-wide 5-hmC in ASD, although recent studies from our group quantified levels of 5-hmC across the genome in human cortex and cerebellum^55^ and across neurodevelopment^56^.

To conclude, we identified widespread differences in DNA methylation associated with iASD, with consistent signals in both cortical regions that were distinct to those observed in the cerebellum. Individuals carrying a duplication on chromosome 15q (dup15q), representing a genetically-defined subtype of ASD, were characterized by striking differences in DNA methylation across a discrete domain spanning an imprinted gene cluster within the duplicated region. In addition to the striking cis-effects on DNA methylation observed in dup15q carriers, we identified convergent methylomic signatures associated with both iASD and dup15q, reflecting the findings from previous studies of gene expression and H3K27ac. Our study represents the first systematic analysis of DNA methylation associated with ASD across multiple brain regions, providing novel evidence for convergent molecular signatures associated with both idiopathic and syndromic autism and highlighting potential disease-associated pathways that warrant further investigation.

## Materials and methods

### Post-mortem brain tissue from autism cases and controls

Tissue samples for this study were acquired from the Autism Tissue Program (ATP) brain bank at the Oxford UK Brain Bank for Autism (www.brainbankforautism.org.uk), Harvard Brain and Tissue Bank (https://hbtrc.mclean.harvard.edu) and the National Institute for Child Health and Human Development (NICHD) Eunice Kennedy Shriver Brain and Tissue Bank for Developmental Disorders. (http://www.medschool.umaryland.edu/btbank/). All subjects were de-identified prior to acquisition and all samples were dissected by trained neuropathologists, snap-frozen and stored at −80 °C. A total of 223 samples obtained from 95 individuals were included in this study with up to up to three brain regions from each individual donor: dorsolateral or medial prefrontal cortex (corresponding to Brodmann area 9 (BA9) and denoted as ‘prefrontal cortex’ (PFC)), superior temporal gyrus (corresponding to BA41, BA42, or BA22 and denoted as ‘temporal cortex’ (TC)), and cerebellar vermis (CB) (**Supplementary Figure 1**). Further information about the samples is given in **Supplementary Table 1**. Genomic DNA was isolated from ∼100 mg of each dissected brain region using a standard phenol-chloroform extraction method, and tested for degradation and purity before analysis.

### DNA methylation profiling and data quality control

All samples were randomized with respect to phenotypic status, age, sex and brain-bank to avoid batch effects throughout all experimental procedures. Genomic DNA (500ng) from each sample was treated in duplicate with sodium bisulfite using the Zymo EZ DNA Methylation-Lightning Kit™ (Zymo Research, Irvine, CA, USA). Genome-wide DNA methylation was quantified using the Illumina Infinium HumanMethylation450 BeadChip (“450K array) (Illumina, San Diego, CA, USA) and scanned on an Illumina HiScan System (Illumina, San Diego, CA, USA). Illumina GenomeStudio software (Illumina, San Diego, CA, USA) was used to extract signal intensities for each probe, generating a final report that was imported in to the R statistical environment 3.0.2 (www.r-project.org) ^57^ using the *methylumi* ^58^ package. Data quality control and pre-processing were performed using the *wateRmelon* package as described previously ^59^. Multidimensional scaling plots of sex chromosome probes were used to check that the predicted sex corresponded with the reported sex for each individual and comparison of 65 SNP probes on the array confirmed that matched tissues were sourced from the same individual. Stringent filtering of the pre-normalized Illumina 450K data was performed. First, we removed the 65 SNP probes, cross-reactive probes and probes overlapping polymorphic CpGs containing an SNP with minor allele frequency>5% within 10 bp of the single base extension position as detailed in the Illumina annotation file and identified in recent publications ^60, 61^. Second, CpG sites with a detection p-value of >0.05 in 1% of samples identified by the *pfilter* function within the *wateRmelon* R package were removed. Third, polymorphic single nucleotide polymorphism (SNP) control probes (n=65) located on the array were used to confirm that matched cortex and cerebellum tissues were sourced from the same individual. The final dataset consists of a total of 417,460 probes from 76 PFC samples (n=36 iASD patients, n=7 dup15q patients, n=33 controls), 77 TC samples (n=33 iASD patients, n=6 dup15q patients, n=38 controls) and 70 CB samples (n=34 iASD patients, n=7 dup15q patients, n=29 controls) (**Supplementary Table 1**) was normalized with the *dasen* function of the *wateRmelon* R package and then batch-corrected with the *ComBat* function of the *ComBat* R package ^62^.

### Identification of autism-associated differential methylation

All statistical analyses were conducted using R statistical package (version 3.1.1). Analyses were performed to test for differential methylated positions (DMPs) and differentially methylated regions (DMRs) associated with disease status for each brain region. The R package *Cell EpigenoType Specific* (CETS) mapper ^63^ designed for the quantification and normalisation of differing neuronal proportions in genome-wide DNA methylation data sets was used to estimate brain cellular heterogeneity in cortex (both PFC and TC) samples. CETS based neuronal cell composition estimate was not applied on the cerebellum samples given the known high proportion of non-NeuN-expressing neurons in this brain region ^41, 63^. To model the effect of sample-specific variables, we performed linear regression for each probe using age, gender, brain bank, CETS (for PFC and TC, but not CB) and diagnosis as independent variables. We also applied an LME model framework for samples with data from multiple cortical regions to identify consistent differential ASD-associated DNA methylation markers across the cortex. The individual donor identified was treated as a random effect, and age, gender, brain bank, CETS, brain region and diagnosis were treated as fixed effects. ASD-associated differential methylated regions (DMRs) were identify using the *Python* module *comb-p* ^44^ to group spatially correlated DMPs (seed P-value<1.00E-03, minimum of 2 probes) at a maximum distance of 300 bp in each analytical groups. Significant DMRs were identified as those with at least two probes and a corrected *P* value < 0.05 using Sidak correction ^64^.

### Weighted gene correlation network analysis

For the co-methylation network analyses, modules of co-methylated probes were identified using the WGCNA package in R ^45^ and network was constructed for cross-cortex samples. For each network, the input probe set was pruned to remove probes with minimal variability across samples. We required that each probe have a minimum range of methylation values of 5% within the middle 80% of samples, resulting in 251,311 probes for cortex. In the cross-cortex network, to enrich for modules relating to diagnosis, we regressed out the effect of CET score and age after fitting a linear model for each probe with diagnosis, age, sex, cortical region, CET score, and brain bank as independent variables. A signed network was constructed using the biweight midcorrelations between probes with a soft power of 7. This network was generated blockwise, using the WGCNA function blockwiseModules with the parameters: maxBlockSize=15000, mergeCutHeight=0.1, deepSplit=4, and minModuleSize=100. For each module, a linear mixed effects model was fit between its eigengene and diagnosis, age, sex, cortical region, batch, CET score, and brain bank as fixed effects with individual ID as a random effect. This LME model was fit for 3 subsets of samples: (iASD + dup15q) vs control, iASD vs control, and dup15q vs control.

### Gene ontology pathway analysis

Illumina UCSC gene annotation, which is derived from the genomic overlap of probes with RefSeq genes or up to 1500 bp of the transcription start site of a gene, was used to create a test gene list from the probes identified in the disease-associated modules for pathway analysis. Where probes were not annotated to any gene (i.e. in the case of intergenic locations), they were omitted from this analysis; where probes were annotated to multiple genes, all were included. A logistic regression approach was used to test if genes in this list predicted pathway membership, while controlling for the number of probes that passed quality control (i.e., were tested) annotated to each gene. Pathways were downloaded from the GO website (http://geneontology.org/) and mapped to genes, including all parent ontology terms. All genes with at least one 450 K probe annotated and mapped to at least one GO pathway were considered. Pathways were filtered to those containing between 10 and 2000 genes. After applying this method to all pathways, the list of significant pathways (*P* < 0.05) was refined by grouping to control for the effect of overlapping genes. This was achieved by taking the most significant pathway, and retesting all remaining significant pathways while controlling additionally for the best term. If the test genes no longer predicted the pathway, the term was said to be explained by the more significant pathway, and hence these pathways were grouped together. This algorithm was repeated, taking the next most significant term, until all pathways were considered as the most significant or found to be explained by a more significant term.

## Acknowledgements

This work was funded by PsychENCODE Grant 1R01MH094714 to DHG. JM is supported by Medical Research Council grants MR/R005176/1, MR/K013807/1, and MR/M008924/1. We are grateful to the patients and families who participate in the tissue programs from which our samples are obtained. Human tissue was obtained from the Autism BrainNet (sponsored by the Simons Foundation and Autism Speaks, formerly the Autism Tissue Program) and the University of Maryland Brain and Tissue Bank, which is a component of the NIH NeuroBioBank and the Oxford Brain Bank.

## Conflicts of interest

None of the authors have any conflicts of interest to declare.

